# Development of two microbial source tracking markers for detection of wastewater-associated *Escherichia coli* isolates

**DOI:** 10.1101/2022.08.23.505042

**Authors:** Ryota Gomi, Eiji Haramoto, Hiroyuki Wada, Yoshinori Sugie, Chih-Yu Ma, Sunayana Raya, Bikash Malla, Fumitake Nishimura, Hiroaki Tanaka, Masaru Ihara

## Abstract

*Escherichia coli* has been used as an indicator of fecal pollution in environmental waters. However, its presence in environmental waters does not provide information on the source of water pollution. Identifying the source of water pollution is paramount to be able to effectively reduce contamination. The present study aimed to identify *E. coli* microbial source tracking (MST) markers that can be used to identify domestic wastewater contamination in environmental waters. We first analyzed wastewater *E. coli* genomes sequenced by us (n = 50) and RefSeq animal *E. coli* genomes of fecal origin (n = 82), and identified 144 candidate wastewater-associated marker genes. The sensitivity and specificity of the candidate marker genes were then assessed by screening the genes in 335 RefSeq wastewater *E. coli* genomes and 3,318 RefSeq animal *E. coli* genomes. We finally identified two MST markers, namely W_nqrC and W_clsA_2, which could be used for detection of wastewater-associated *E. coli* isolates. These two markers showed higher performance than the previously developed human wastewater-associated *E. coli* markers H8 and H12. When used in combination, W_nqrC and W_clsA_2 showed specificity of 98.9% and sensitivity of 25.7%. PCR assays to detect W_nqrC and W_clsA_2 were also developed and validated. The developed PCR assays are potentially useful for detecting *E. coli* isolates of wastewater origin in environmental waters, though users should keep in mind that the sensitivity of these markers is not high. Further studies are needed to assess the applicability of the developed markers to a culture-independent approach.

## 1. Introduction

Surface waters contaminated by wastewater can increase human health risks because untreated and insufficiently-treated wastewater contains various pathogens (Castro-Hermida et al., 2008; Naidoo and Olaniran, 2013; Zhi et al., 2020). Humans can be exposed to such pathogens through activities such as swimming and the consumption of foods irrigated with poor-quality water (Boehm et al., 2018; Steele and Odumeru, 2004). Because it is impractical to test for all possible pathogens in surface waters, it is a standard practice to monitor fecal indicator bacteria (FIB), such as total coliforms, fecal coliforms, and *Escherichia coli* (Field and Samadpour, 2007; Naidoo and Olaniran, 2013). Although FIB are not normally pathogenic to humans, their presence in water indicates the possible presence of pathogens (e.g., *Campylobacter* spp., *Salmonella* spp., and *Cryptosporidium parvum*), which also live in human and animal digestive systems (Ahmed et al., 2019b). FIB can originate from many sources, such as community sewage (containing human-specific pathogens and zoonotic pathogens) and diffuse pollution from animal farms and wildlife (containing zoonotic pathogens). The major limitation of FIB is that, due to their cosmopolitan nature, their presence does not provide information on the possible origin of contamination (Warish et al., 2015). Identifying the origin of water pollution is paramount in order to assess the associated health risks and determine the actions needed to reduce contamination (Scott et al., 2002).

Microbial source tracking (MST) is an approach used to determine the sources of fecal contamination in environmental waters (Harwood et al., 2014). Because culture-independent methods have many advantages, such as being fast and not labor-intensive, the majority of MST methods in use today are culture-independent. For example, markers such as the 16S rRNA gene of *Bacteroides* spp. and pepper mild mottle virus (PMMoV) have been used to track the sources of fecal contamination in MST in a culture-independent manner (Gonzalez-Fernandez et al., 2021; Mathai et al., 2020). However, routine water quality assessment has focused primarily on traditional FIB, and alternative indicators such as *Bacteroides* spp. are not usually monitored for this purpose (Senkbeil et al., 2019; Teixeira et al., 2020). Because *E. coli* is still a widely-used FIB, MST markers targeting *E. coli* would be useful if included in the course of routine monitoring. Even a culture-dependent approach would be useful for *E. coli*, because users do not have to collect isolates exclusively for MST purposes but can use *E. coli* colonies obtained in routine water quality assessments.

*E. coli* has been used and studied as an indicator of fecal contamination for many years. Some previous studies have focused on host-specific traits of this organism to identify potential sources of fecal contamination. Earlier studies employed library-dependent techniques such as rep-PCR, ribotyping, and denaturing gradient gel electrophoresis (DGGE) to differentiate *E. coli* from different sources (Buchan et al., 2001; Carson et al., 2001; Dombek et al., 2000). Later, a few studies identified library-independent *E. coli* markers such as cattle and swine-specific *E. coli* toxin genes (Khatib et al., 2002; 2003), though none seem to be widely used today. While *E. coli* has been used in both library-dependent and -independent MST studies, it has been suggested that *E. coli* is not an ideal target for MST studies because it may not be distinct enough to be separated into host-specific groups (Ishii and Sadowsky, 2008). The sharing/transmission of *E. coli* clones among different host species has also been reported (Johnson and Clabots, 2006; Li et al., 2019; Stenske et al., 2009). However, a number of studies, for example those based on genome analysis, also report certain host-associated traits of *E. coli* (Lupolova et al., 2017; Yu et al., 2021), indicating there to be some level of host-specificity in this species. Although little success has been achieved in developing MST markers targeting

*E. coli*, we have previously identified a set of human wastewater, cattle, pig, and chicken feces-associated *E. coli* MST markers (Gomi et al., 2014). The following studies reported two of the proposed markers, namely H8 and H12, showed relatively high sensitivity and specificity and could be useful as human wastewater-associated markers (Nopprapun et al., 2020; Senkbeil et al., 2019; Warish et al., 2015). However, the original genetic markers were developed by analyzing a small number of *E. coli* genomes (n = 22). Since then, a growing number of *E. coli* genomes have become available in public databases. These *E. coli* genomes include those from various isolation sources and geographically diverse hosts, which might allow the development of reliable MST markers that can be used globally. Here, we analyzed *E. coli* genomes in the database of the National Center for Biotechnology Information (NCBI) as well as genomes sequenced by us to identify more reliable *E. coli* genetic markers for tracking domestic wastewater contamination in environmental waters. Our aim in the present study was to identify culture-dependent MST markers, but applicability to wastewater samples was also tested in a culture-independent manner. The sensitivity and specificity of the previously developed human wastewater-associated markers H8 and H12 were also re-evaluated.

## 2. Materials and methods

### 2.1. *E. coli* isolation from wastewater and genome sequencing

The flowchart of the study design is shown in **Figure 1**. We obtained wastewater *E. coli* isolates from three municipal wastewater treatment plants (WWTPs), namely WWTP A, WWTP B, and WWTP C, in Shiga Prefecture, Japan. The amount of sewage treated by each WWTP per day is 80,000 m^3^ for WWTP A, 60,000 m^3^ for WWTP B, and 250,000 m^3^ for WWTP C. Almost all the areas served by these WWTPs have separate sewer systems, and the WWTPs mainly receive domestic wastewater. Biologically treated wastewater samples before/after chlorination were collected from the WWTPs between December 2016 and December 2017. The chlorinated samples were treated with sodium thiosulfate immediately after collection to neutralize the chlorine. Samples were collected using sterile sampling bags or sterile sampling bottles, stored at 4 °C, and processed within 24 hours. Samples were serially diluted with PBS (ten-fold serial dilution), and 1 mL of undiluted/diluted samples were processed using the pour plate method with XM-G agar (Nissui, Tokyo, Japan). Plates were incubated at 37 °C for 18h, and colonies were randomly picked from the plates. Colonies suspected to be mixed were sub-cultured with fresh XM-G agar plates to obtain pure isolates. A total of 50 isolates (29 isolates from WWTP A, nine isolates from WWTP B, and 12 isolates from WWTP C) were obtained for genome sequencing (see **Table S1** for information, such as sample type and collection date, on the collected *E. coli*).

**Figure 1.**
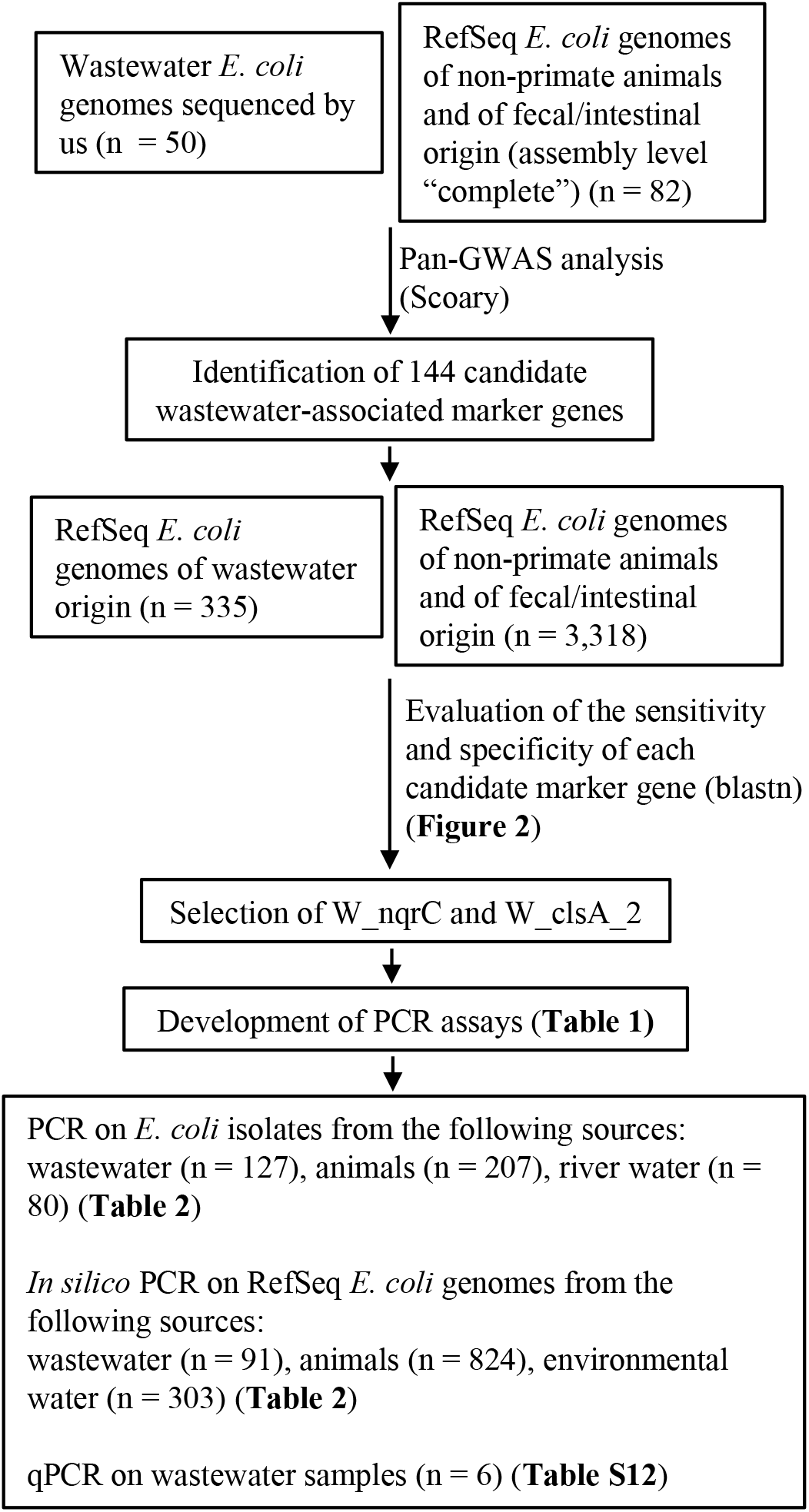
Overview of the strategy to identify and test the wastewater-associated *E. coli* genetic markers. RefSeq *E. coli* genomes used for blastn analysis (335 wastewater *E. coli* genomes and 3,318 animal *E. coli* genomes) and *in silico* PCR analysis (91 wastewater *E. coli* genomes, 824 animal *E. coli* genomes, and 303 environmental *E. coli* genomes) included both complete and draft genome assemblies. There was no overlap between the genomes used for blastn analysis and *in silico* PCR analysis.

Colonies were resuspended in PBS and centrifuged to prepare a cell pellet. DNA was extracted from the pellet using a DNeasy blood and tissue kit (Qiagen, Hilden, Germany) following the manufacturer’s instructions, and the concentration of extracted DNA was measured using a Qubit fluorometer with dsDNA High sensitivity kit (Thermo Fisher Scientific, Waltham, MA, United States). Extracted DNA was stored at -30°C for up to a week until sample preparation for genome sequencing. The extracted DNA was processed using a Nextera XT DNA sample preparation kit (Illumina, San Diego, CA) and sequenced on an Illumina MiSeq instrument for 600 cycles (300-bp paired-end sequencing) as described previously (Ma et al., 2022).

### 2.2. Retrieving complete *E. coli* genome sequences from the NCBI database

The Reference Sequence (RefSeq) *E. coli* genomes with an assembly level of “complete genome” (n = 1,278) were downloaded in May 2021. Metadata information, such as strain name, host, and isolation source, was extracted from the GenBank files using in-house Python scripts, which uses tools such as Biopython Bio.SeqIO module (Cock et al., 2009). *E. coli* genomes of non-primate animals and of fecal/intestinal origin (n = 82) were kept for further analysis (*E. coli* genomes from birds were also included here. See **Table S2** for information on these animal *E. coli* genomes).

### 2.3. Genome assembly and annotation

Raw sequence reads from wastewater *E. coli* were trimmed using fastp (v0.20.0) (Chen et al., 2018), and the trimmed reads were assembled using Unicycler (v0.4.8) with the --no_correct option (Wick et al., 2017). For the complete RefSeq genomes, we generated simulated paired-end reads using wgsim (v1.12) (https://github.com/lh3/wgsim). Simulated reads were trimmed using fastp and assembled using Unicycler with the same parameters as those used for assembling the wastewater *E. coli* genomes. We did not use the complete RefSeq genomes but used these draft assemblies in the following analysis. This was to remove any biases introduced by differences in assembly status (i.e., complete assemblies vs. draft assemblies) in the pan-genome-wide association study (pan-GWAS). Assembled genomes were annotated using Prokka (v1.14.6) (Seemann, 2014).

### 2.4. Identification of candidate wastewater-associated marker genes using a pan-GWAS approach

The pan-genome of 132 *E. coli* genomes (50 wastewater *E. coli* genomes + 82 animal *E. coli* genomes) was constructed by running Roary (v3.13.0) on GFF3 files generated by Prokka (Page et al., 2015). The -s option was used to turn off paralog splitting. The pan-GWAS approach implemented by Scoary was used to find genes that were enriched in wastewater *E. coli* genomes (Brynildsrud et al., 2016). Scoary (v 1.6.16) was run on Roary’s gene presence/absence file. The --collapse option was used to collapse genes that had identical distribution patterns among the genomes. Because our aim was to identify genes that were enriched in wastewater *E. coli* genomes and not to infer causal association, the --no_pairwise option was used as recommended by the developers (https://github.com/AdmiralenOla/Scoary). Genes with Benjamini-Hochberg adjusted p-values of lower than 0.05 and with a specificity exceeding 95% were defined as candidate wastewater-associated marker genes. We set the specificity threshold so as to reduce the number of false positives, considering the future application to DNA directly extracted from environmental samples (this type of DNA can contain DNA from multiple *E. coli* isolates and thus is sensitive to false positives). When a set of collapsed genes met the p-value and specificity criteria, one gene was randomly selected as a candidate marker.

### 2.5. Retrieving genome sequences from the NCBI database for sensitivity and specificity analysis

The RefSeq *E. coli* genomes (n = 20,237), including both complete and draft assemblies, were downloaded in June 2021. Metadata information was extracted from the GenBank files using in-house Python scripts. Genomes of wastewater origin (n = 335) and genomes of non-primate animals and of fecal/intestinal origin (n = 3,318) were kept for sensitivity and specificity analysis (see **Table S3** for information on the wastewater *E. coli* genomes and **Table S4** for information on the animal *E. coli* genomes).

### 2.6. Evaluating the sensitivity and specificity of candidate marker genes

The candidate marker genes were detected in 3,653 genome assemblies (335 RefSeq wastewater *E. coli* genomes + 3,318 RefSeq animal *E. coli* genomes) using the blastn program with the default parameters. Thresholds of a minimum percent identity of 80 % and a minimum DNA coverage of 50 % were used to define the presence of a gene. For each candidate marker gene, the following metrics were calculated: true positive (TP) (the number of wastewater *E. coli* genomes positive for the candidate marker), true negative (TN) (the number of animal *E. coli* genomes negative for the candidate marker), false positive (FP) (the number of animal *E. coli* genomes positive for the candidate marker), and false negative (FN) (the number of wastewater *E. coli* genomes negative for the candidate marker). Sensitivity [TP/(TP + FN)] and specificity [TN/(TN + FP)] were then calculated for each candidate marker gene (note that sensitivity and specificity were reported as %). The above blastn analysis and calculation were also performed for H8 and H12. Note that H8 corresponded to one of the candidate marker genes.

### 2.7. Development of PCR assays

Primers that yield desired product sizes were designed for the identified marker genes as shown in **Table 1**. For each target gene, the validity of the primers was checked with Primer 3 software using the default settings (Untergasser et al., 2012). Moreover, using Primer-BLAST with the default settings, we confirmed that an amplicon of the desired size was produced from each wastewater genome with the target gene that had been sequenced in **2.1** (Ye et al., 2012).

**Table 1.**
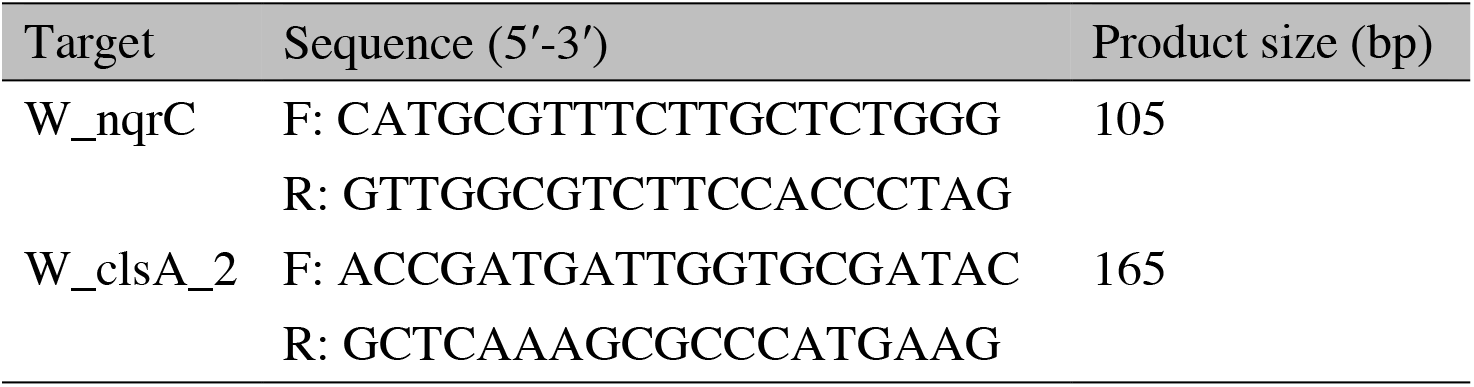
Primers designed in this study.

To further validate the developed PCR primers, we performed PCR on *E. coli* isolates collected from the following sources: wastewater influent (n = 78), wastewater effluent (after biological treatment and chlorination) (n = 49), cattle (n = 39), chickens (n = 30), deer (n = 30), pigs (n = 30), waterfowl (n = 30), cats (n = 24), and dogs (n = 24). Detailed information on these isolates is available in **Table S5**. Briefly, wastewater influent isolates (n = 48) and effluent isolates (n = 49) were newly collected in 2021 from WWTP A in Shiga Prefecture, Japan, and other wastewater influent isolates (n = 30) were collected in 2018 from a WWTP (WWTP D) in Yamanashi Prefecture, Japan. WWTP D treats 110,000 m^3^ of sewage per day and the area served by this WWTP has separate sewer systems. We used the same sampling protocol and *E. coli* isolation procedure as described in **2.1**, except that we used Chromocult coliform agar (Merck, Darmstadt, Germany) or CHROMagar ECC (CHROMagar, Paris, France) and incubated the plates for ∼24 hours at 37 °C to isolate *E. coli* from the influent samples collected from WWTP D. Animal *E. coli* isolates were obtained from animal feces collected in Kyoto Prefecture, Nara Prefecture, Shiga Prefecture, or Yamanashi Prefecture between 2011 and 2022, as described in **Table S5**. All animal feces were collected using sterile sampling bags or sterile centrifuge tubes, stored at 4 °C, and processed within 24-48 hours. Animal feces were directly streaked onto XM-G agar using sterile swabs/inoculating needles or suspended and diluted with PBS and processed using the pour plate method with XM-G agar (for two deer fecal samples, the pour plate method with CHROMagar ECC was used). The plates were incubated for 18-24 hours at 37 °C. Sub-culturing was performed to obtain pure colonies. Up to three isolates were obtained from a single individual (Dombek et al., 2000).

DNA was extracted from *E. coli* obtained from WWTP A and cattle and deer feces using a DNeasy blood and tissue kit. The concentration of extracted DNA was measured using a Qubit fluorometer with dsDNA High sensitivity kit. The extracted DNA was stored at -30°C for up to one year until the PCR experiments. For *E. coli* obtained from WWTP D and other animal feces, we used cell suspensions as the inputs in the PCR experiments. Cell suspensions were prepared by either of the following two methods: (i) a single *E. coli* colony grown on the XM-G plate was picked up using a clean tip, suspended in 500 μL of ultrapure water, and then vortexed; or (ii) a glycerol stock (*E. coli* stored in 25% glycerol) was diluted 10 times using ultrapure water. PCR reactions were performed on the extracted DNA or cell suspension using QuantiFast SYBR Green PCR Kit (Qiagen, Hilden, Germany) or QuantiNova SYBR Green PCR Kit (Qiagen) with PCR conditions as described below. For QuantiFast SYBR Green PCR Kit, PCR mixture (25 μL) was composed of 12.5 μL of 2× QuantiFast SYBR Green PCR Master Mix, 2.5 μL each of forward and reverse primers (5 μM), 5 μL of template DNA or cell suspension, and 2.5 μL of PCR grade water or ultrapure water. The reactions were carried out by incubation at 95 °C for 5 min, followed by 25 to 30 cycles (depending on the template concentrations) consisting of 95 °C (denaturation) for 10 s and 60 °C (combined annealing/extension) for 30 s. A melting curve analysis was performed after the amplification. For QuantiNova SYBR Green PCR Kit, PCR mixture (20 μL) was composed of 10 μL of 2× QuantiNova SYBR Green PCR Master Mix, 2.8 μL each of forward and reverse primers (5 μM), 2 μL of template DNA or cell suspension, and 2.4 μL of PCR grade water or ultrapure water. The reactions were carried out by incubation at 95 °C for 2 min, followed by 30 cycles consisting of 95 °C (denaturation) for 5 s and 60 °C (combined annealing/extension) for 10 s.

A melting curve analysis was performed after the amplification. In addition to the primers for W_nqrC and W_clsA_2, we also used previously published primers for amplifying H8, H12, and also *uidA* (primers UAL and UAR): we included *uidA* primers to confirm the *E. coli* identification (Gomi et al., 2014; Maheux et al., 2009). All PCR reactions were performed on a Thermal Cycler Dice Real Time System III (Takara Bio, Inc.). Previously sequenced *E. coli* isolates, namely KOr026 (positive: W_clsA_2 and H8, negative: W_nqrC and H12, BioSample accession: SAMD00053091) and KMi027 (positive: W_nqrC and H12, negative: W_clsA_2 and H8, BioSample accession: SAMD00053165), were used as positive/negative controls for both extracted DNA and cell suspensions (Gomi et al., 2017b). We also included no template controls. The correct PCR products were confirmed by comparison with the melting curves of the positive controls, which showed peaks at melting temperatures of 82.5 °C, 89.0 °C, 91.6 °C, 85.7 °C, and 84.4 °C for W_nqrC, W_clsA_2, H8, H12, and *uidA*, respectively. We performed replicate reactions (duplicate reactions) for a subset of samples (50 samples, including 42 for extracted DNA and 8 for cell suspensions) and confirmed the results to be consistent between all technical replicates.

We also performed *in silico* PCR using the developed primers on RefSeq *E. coli* genomes that were made public after June 2021 (i.e., genomes that were not used for blastn analysis). This was to test the developed PCR primers using *E. coli* genomes of diverse origins and from various geographical locations. *In silico* PCR was previously shown to be highly concordant with *in vitro* PCR when used for gene detection in *E. coli* (Beghain et al., 2018; Lindsey et al., 2017). Briefly, RefSeq *E. coli* genomes, including both complete and draft assemblies, were downloaded in August 2022. Among the downloaded genomes, those made public after June 2021 were retained (n = 6,399). Metadata information was extracted from the retained GenBank files, and the genomes of wastewater origin (n = 91) and those of non-primate animals and of fecal/intestinal origin (n = 824) were retrieved. *In silico* PCR was performed on the Primer-BLAST website using primer sequences of W_nqrC, W_clsA_2, H8, and H12 with the default parameters, except that we used 500 bp as max target amplicon size (Ye et al., 2012).

### 2.8. Detection of MST markers in *E. coli* isolates from environmental waters

*E. coli* isolates were obtained in January 2022 from a river in Kyoto Prefecture in Japan. The sampling points in the river were downstream of two wastewater treatment plants and known to be heavily affected by treated wastewater (**Figure S1**). A total of eight water samples were collected from the river using sterile sampling bags, stored at 4 °C, and processed within 24 hours. Ten *E. coli* colonies were obtained from each sample using the membrane filter method with XM-G agar plates. The plates were incubated for 18 hours at 37 °C, and colonies were sub-cultured with fresh XM-G agar plates to obtain pure isolates. PCR reactions were performed using cell suspensions as described above to test for the presence of W_nqrC and W_clsA_2 in the obtained *E. coli* isolates.

We also performed *in silico* PCR on RefSeq *E. coli* genomes of environmental water origin. Environmental *E. coli* genomes were extracted from the RefSeq *E. coli* genomes downloaded in August 2022 based on the metadata information. *In silico* PCR was performed using the retrieved genomes (n = 303) on the Primer-BLAST website as described above.

### 2.9. Quantitative PCR (qPCR) assays on DNA extracts from wastewater samples

qPCR assays were performed to determine the concentrations of the developed markers in total DNA extracted from wastewater samples. Three primary effluent samples and three biologically treated wastewater samples before chlorination were taken from WWTP A on different occasions in October and November 2022. Primary effluent samples (40–80 mL) and treated wastewater samples (1000 mL) were filtered through 0.2 μm pore size polycarbonate membranes (Advantec, Tokyo, Japan). Multiple membranes were used when membranes became clogged during filtration. Filters were stored at -30°C until DNA extraction. DNA was extracted from each filter using a FastDNA SPIN Kit for Soil (MP Biomedicals, Irvine, CA), and PCR inhibitors were removed from the DNA extracts using CHROMA SPIN+TE-400 Columns (Takara Bio, Inc.) and Amicon Ultra-0.5 (100K) Centrifugal Filter Devices (Millipore, Darmstadt, Germany). qPCR was performed using a QuantiNova SYBR Green PCR Kit under the same conditions as PCR performed on *E. coli* isolates, except that 35 cycles of reactions were performed. The qPCR reactions were performed in duplicate. The correct PCR products were confirmed by comparison with the melting curves of the positive controls. qPCR standards were prepared using 10-fold serial dilutions of genomic DNA extracted from pure cultures of KOr026 and KMi027. The copy numbers (T) of genomic DNA used for standard curve construction were estimated using the equation T = [D/(5 × 10^6^ × 660)] × 6.022 × 10^23^, where D (g/μl) is the DNA concentration.

### 2.10. Sequence Data Accession Number(s)

The raw sequencing data for the wastewater *E. coli* isolates were deposited in the NCBI SRA under BioProject accession number PRJNA765288.

## 3. Results and discussion

### 3.1. Basic characteristics of wastewater *E. coli* genomes sequenced by us

The 50 wastewater *E. coli* genomes sequenced by us were highly diverse, comprising 44 different sequence types (STs). These draft assemblies had a median of 140 contigs (range 49 contigs to 346 contigs), a median genome size of 4,875,061 bp (range 4,576,995 bp to 5,375,884 bp), and a median N50 of 179,046 bp (range 63,435 bp to 737,274 bp) after removing contigs shorter than 100 bp (**Table S1**). All 50 assemblies met the quality control criteria of (i) contig numbers: ≤ 800; (ii) genome size: 3.7 Mbp to 6.4 Mbp; and (iii) N50: >20 kb, as recommended by EnteroBase (Zhou et al., 2020).

### 3.2. Identification of candidate wastewater-associated marker genes

A pan-genome was constructed with Roary using 50 wastewater *E. coli* genomes sequenced by us and 82 RefSeq animal *E. coli* genomes. Roary identified a total of 25,343 genes, including 2,925 core genes (genes shared among ≥ 99% of strains) and 22,418 accessory genes (genes observed among < 99% of strains). The pan-GWAS approach was employed to identify candidate wastewater-associated genes among the accessory genes. A total of 144 genes met the criteria of Benjamini-Hochberg adjusted p-values of less than 0.05 and specificity higher than 95%, and thus were defined as candidate marker genes (see **Table S6** for details of the identified genes). These 144 genes were subjected to sensitivity and specificity analysis by blastn using a larger number of RefSeq *E. coli* genomes.

### 3.3. Sensitivity and specificity of candidate marker genes determined by blastn analysis

The sensitivity and specificity of 144 candidate marker genes were assessed by screening these genes in 335 RefSeq wastewater *E. coli* genomes and 3,318 RefSeq animal *E. coli* genomes. **Figure 2** shows the distribution of sensitivity and specificity of each candidate marker gene. Most candidate markers achieved high specificity, which is congruent with the fact that we used a threshold of 95% specificity for identifying candidate markers in the Scoary analysis. However, there were no genes that simultaneously showed high sensitivity and high specificity, which may be partially due to the presence of *E. coli* isolates that can colonize multiple host species (Johnson and Clabots, 2006; Stenske et al., 2009). In fact, among the genes showing >70% specificity, no genes showed >40% sensitivity. It should be noted that there were genes with almost 100% sensitivity and 0% specificity (i.e., detected in almost all the tested isolates) even though we selected genes with specificity higher than 95% in the Scoary analysis. This can be because we used blastp with a minimum percent identity of 95% for clustering sequences in the Roary analysis but we used blastn with a minimum percent identify of 80% for gene screening. In other words, genes assigned to different clusters by Roary can share >80% nucleotide sequence identity and thus can be detected in the same blastn analysis. Because the species composition of the hosts of RefSeq animal *E. coli* genomes was biased (some host species were more prevalent than others), the specificity of candidate markers was separately calculated for different host groups, namely, “Cattle”, “Swine”, “Chicken”, “Wild boar (*Sus scrofa*)”, “Canine”, “Sheep and goat”, and “Others” (**Figure S2**). We also considered the specificity calculated for these host groups when selecting MST markers from the candidates.

**Figure 2.**
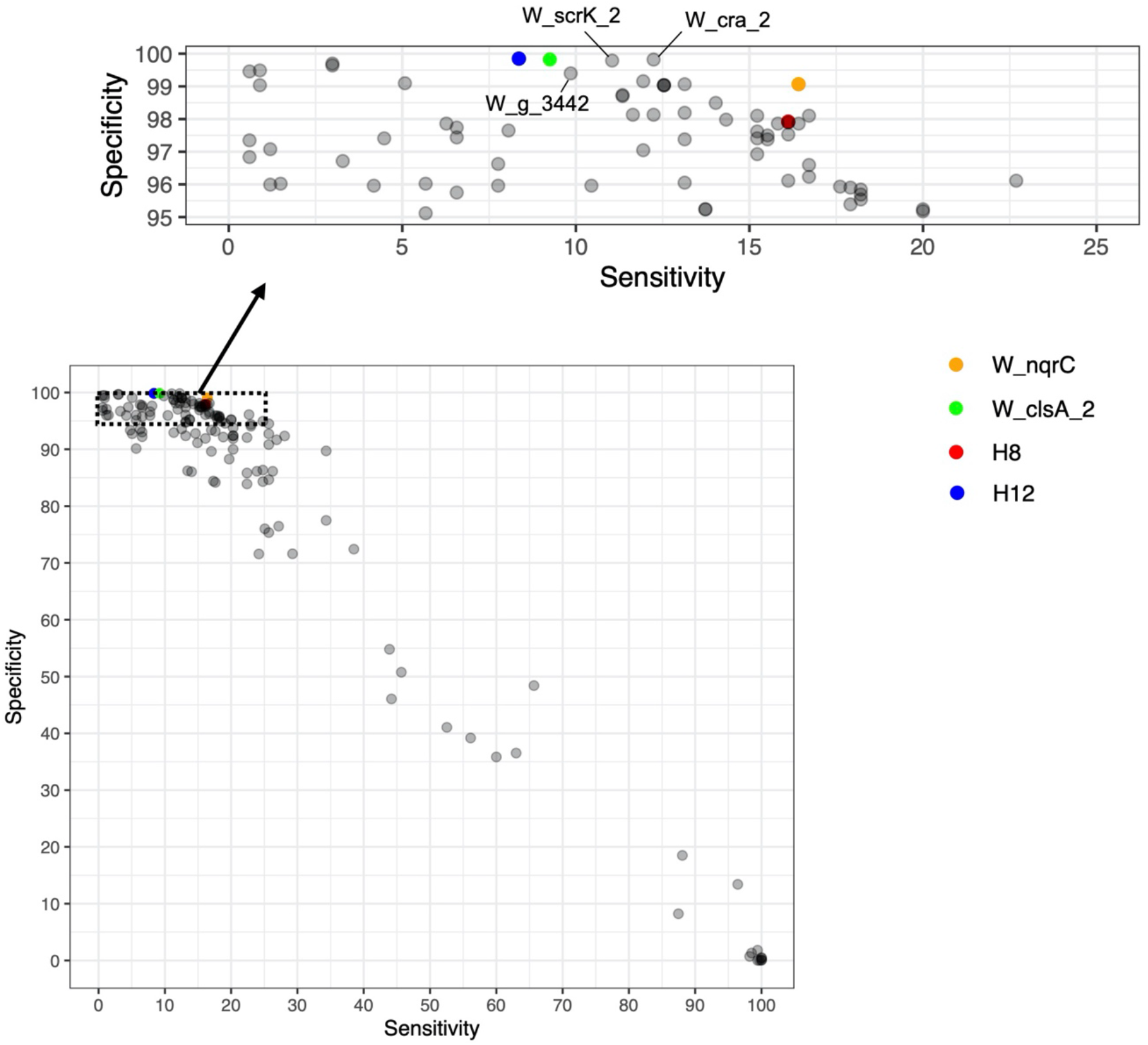
Sensitivity and specificity of candidate wastewater-associated marker genes as determined by blastn analysis. The distribution of sensitivity and specificity of 144 candidate marker genes and previously identified marker genes, H8 and H12, are plotted (H8 in fact corresponded to one of the 144 candidate marker genes). The positions of markers mentioned in the main text (W_scrK_2, W_cra_2, and W_g_3442) are also indicated.

### 3.4. Selection of W_nqrC and W_clsA_2

As noted in the Materials and Methods section, selecting a marker with high specificity is required to reduce the number of false positives, especially when the marker is applied to DNA directly extracted from environmental samples. We therefore selected W_nqrC from the 144 candidates, as this marker showed high specificity (99.1%) and the highest sensitivity (16.4%) among the markers with specificity higher than 99% (**Figure 2**). Moreover, W_nqrC showed the highest sensitivity among the candidate markers which showed >95% specificity for all host groups (**Figure S2**). We then tried to select another marker that was prevalent among the RefSeq wastewater *E. coli* genomes without W_nqrC. This was to increase the sensitivity when both markers are used in combination. Among the candidate markers showing >99% specificity, W_cra_2, W_scrK_2, W_clsA_2, and W_g_3442 showed different marker distribution patterns (the red bars in **Figure S3**) from W_nqrC and also showed relatively high sensitivities (**Figure S3**). Although W_cra_2 and W_scrK_2 showed high specificity (>99.7%), similar sequences with 60%–75% nucleotide sequence identity were prevalent among animal *E. coli* isolates. We therefore selected W_clsA_2 as the second marker because there were no similar sequences prevalent among animal *E. coli*, and this marker showed higher specificity than W_g_3442 (99.8% vs. 99.4%). Moreover, the specificity of W_clsA_2 was 100% for all host groups except “Others” (only six *E. coli* genomes from silver gulls were positive for W_clsA_2) (**Figure S2)**. The sequences of W_nqrC and W_clsA_2 are shown in **Figure S4**. Prokka annotation indicated W_nqrC encodes Na(+)-translocating NADH-quinone reductase subunit C, and W_clsA_2 encodes major cardiolipin synthase ClsA. The BLASTN searches against the nucleotide collection (nr/nt) database revealed that W_nqrC is mostly present on plasmids, while W_clsA_2 is equally present on chromosomes and plasmids. As mentioned in **2.4**., the pan-GWAS approach used in this study simply identifies genes enriched in wastewater *E. coli*, but does not infer causal association. Therefore, W_nqrC and W_clsA_2 may not directly confer adaptive benefits to wastewater *E. coli*. However, these two markers are likely plasmid-associated, which may partially explain the enrichment of these markers in wastewater *E. coli*. This is because genes that confer resistance to antibiotics and heavy metals are also often located on plasmids, which can favor the selection of bacteria with these genes in wastewater (Di Cesare et al., 2016; Karkman et al., 2018; Li et al., 2015). Among the RefSeq wastewater *E. coli* genomes positive for W_nqrC (n = 55), almost half of the genomes (n = 27) were assigned to ST131, which was previously reported to be present among wastewater *E. coli* (Dolejska et al., 2011; Gomi et al., 2017a; Sghaier et al., 2019) and also is known for its association with extraintestinal infections (Nicolas-Chanoine et al., 2014; Riley, 2014; Stoesser et al., 2016). The remaining W_nqrC-positive genomes were assigned to various STs, including ST69 (n = 4), ST95 (n = 4), and ST10 (n = 3). The W_nqrC-positive wastewater *E. coli* genomes sequenced by us (n = 9) belonged to diverse STs, including ST73 (n = 2), ST95 (n = 2), and ST357 (n = 2) (**Table S7**). Among the RefSeq wastewater *E. coli* genomes positive for W_clsA_2 (n = 31), more than half of the genomes (n = 17) were assigned to ST635, which was previously reported to be prevalent among wastewater *E. coli* isolates in Canada and hospital sink *E. coli* isolates in the UK (Constantinides et al., 2020; Zhi et al., 2019). The remaining W_clsA_2-positive genomes belonged to various STs, including ST399 (n = 3), ST401 (n = 2), ST607 (n = 2), and ST3168 (n = 2). The W_clsA_2-positive wastewater *E. coli* genomes sequenced by us (n = 5) belonged to five different STs (ST399, ST472, ST607, ST635, and ST5295) (**Table S8**).

We note that W_nqrC/W_clsA_2-positive RefSeq wastewater *E. coli* genomes originated from various countries, including the Czech Republic, Germany, Canada, the USA, and South Africa (**Table S7, Table S8**). In fact, 43 (78%) of W_nqrC-positive RefSeq wastewater *E. coli* genomes (n = 55) and 31 (100%) of W_clsA_2-positive RefSeq wastewater *E. coli* genomes (n = 31) originated from countries other than Japan, which indicates that these markers may be applicable to regions outside of our study site.

### 3.5. Sensitivity and specificity of H8 and H12

We also assessed the sensitivity and specificity of the two previously developed markers, H8 and H12, which were calculated based on the results of blastn analysis (**Figure 2**). Notably, H8 corresponded to one of the 144 candidate marker genes identified by Scoary. **Figure 2** shows that both H8 and H12 achieved high specificity (97.9% and 99.8%, respectively), which is consistent with previous studies (Senkbeil et al., 2019; Warish et al., 2015). However, the markers proposed in the present study showed better performances. W_nqrC showed higher specificity and sensitivity than H8 (99.1% vs. 97.9% and 16.4% vs. 16.1%, respectively), and W_clsA_2 showed almost the same specificity (∼99.8%) but higher sensitivity than H12 (9.3% vs. 8.4%). Moreover, when used in combination, a combination of W_nqrC and W_clsA_2 showed higher specificity and sensitivity than a combination of H8 and H12 (98.9% vs. 97.8% and 25.7% vs. 24.2%, respectively). These results suggest that although H8 and H12 could be useful for tracking wastewater contamination, W_nqrC and W_clsA_2 may yield better performances.

### 3.6. Development of PCR assays targeting W_nqrC and W_clsA_2

We designed PCR primers for detection of W_nqrC and W_clsA_2. PCR assays on positive controls yielded the expected DNA bands for both W_nqrC (105 bp) and W_clsA_2 (165 bp), whereas no bands were observed for the negative controls (**Figure S5**). The developed PCR assays were applied to detect the MST markers in *E. coli* isolates obtained from wastewater and animal feces (**Table 2**). W_nqrC or W_clsA_2 was detected in 21 (26.9%) and 9 (18.4%) of wastewater influent and effluent isolates, respectively (**Table 2**). Importantly, there was no overlap between the isolates positive for W_nqrC and W_clsA_2, which suggests that the use of W_nqrC and W_clsA_2 in combination could increase the sensitivity. On the other hand, only seven animal *E. coli* isolates (3.4%, six from dogs and one from waterfowl) were positive for W_nqrC or W_clsA_2. We also performed PCR to detect H8 and H12 using the same set of *E. coli* isolates. H8 or H12 was detected in 11 (14.1%) and 7 (14.3%) of wastewater influent and effluent isolates, respectively (**Table 2**). Only two animal *E. coli* isolates (1.0%, one from pigs and one from waterfowl) were positive for H8 or H12. In our PCR experiments, the sensitivities of the W_nqrC and W_clsA_2 combination were higher than those of the H8 and H12 combination for wastewater influent (26.9% vs 14.1%) and wastewater effluent (18.4% vs 14.3%), while the specificity of the W_nqrC and W_clsA_2 combination was lower than that of the H8 and H12 combination (96.6% vs 99.0%). The lower specificity of the W_nqrC and W_clsA_2 combination was attributed to six dog isolates positive for W_nqrC (we note that these six isolates were from two fecal samples, namely three from one and three from the other). The blastn analysis also showed relatively lower specificity of W_nqrC for canine *E. coli* isolates (**Figure S2E**). One possible reason for this is within-household transmission of *E. coli* between dogs and humans (Johnson and Clabots, 2006; Stenske et al., 2009).

**Table 2.**
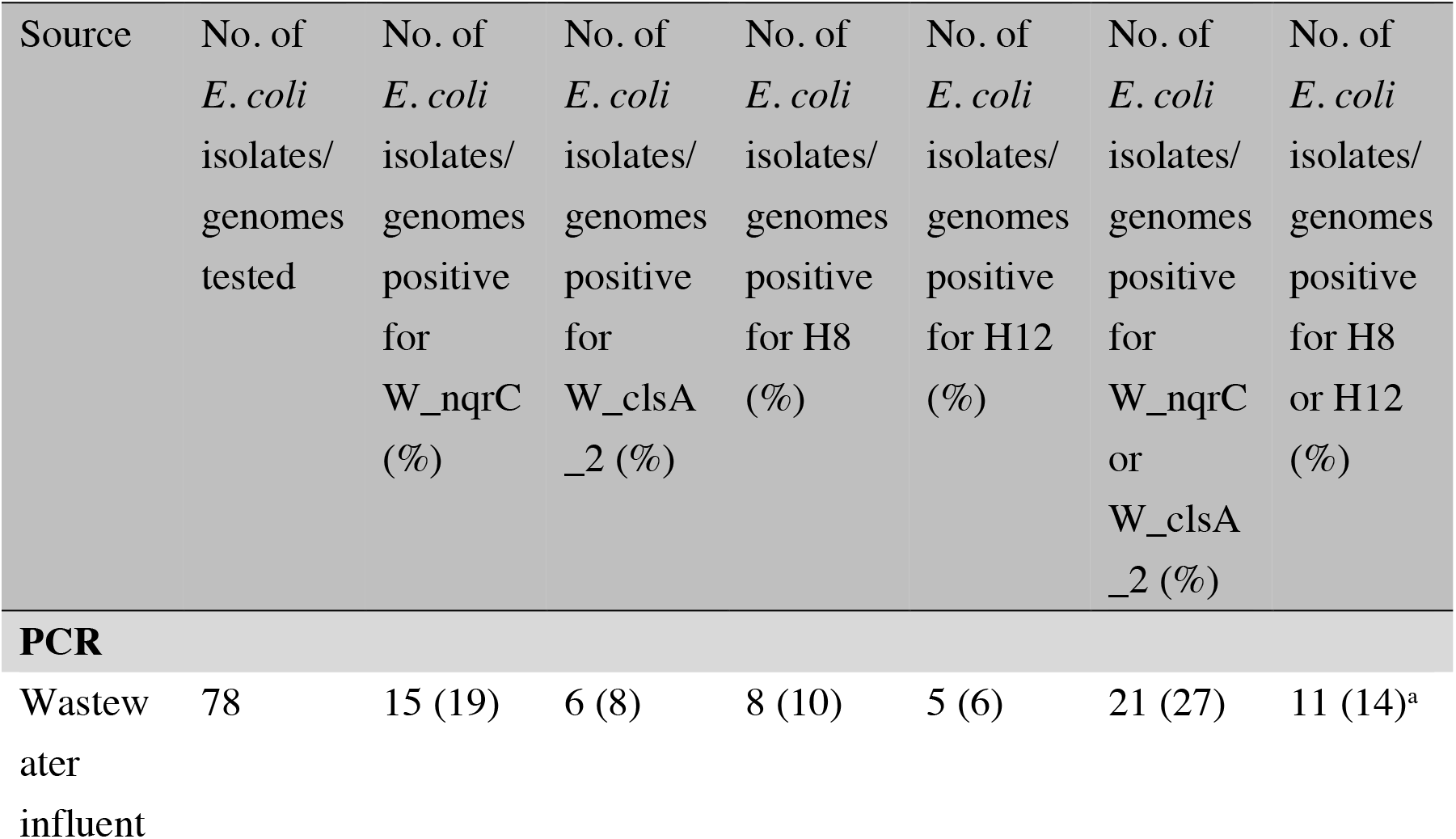
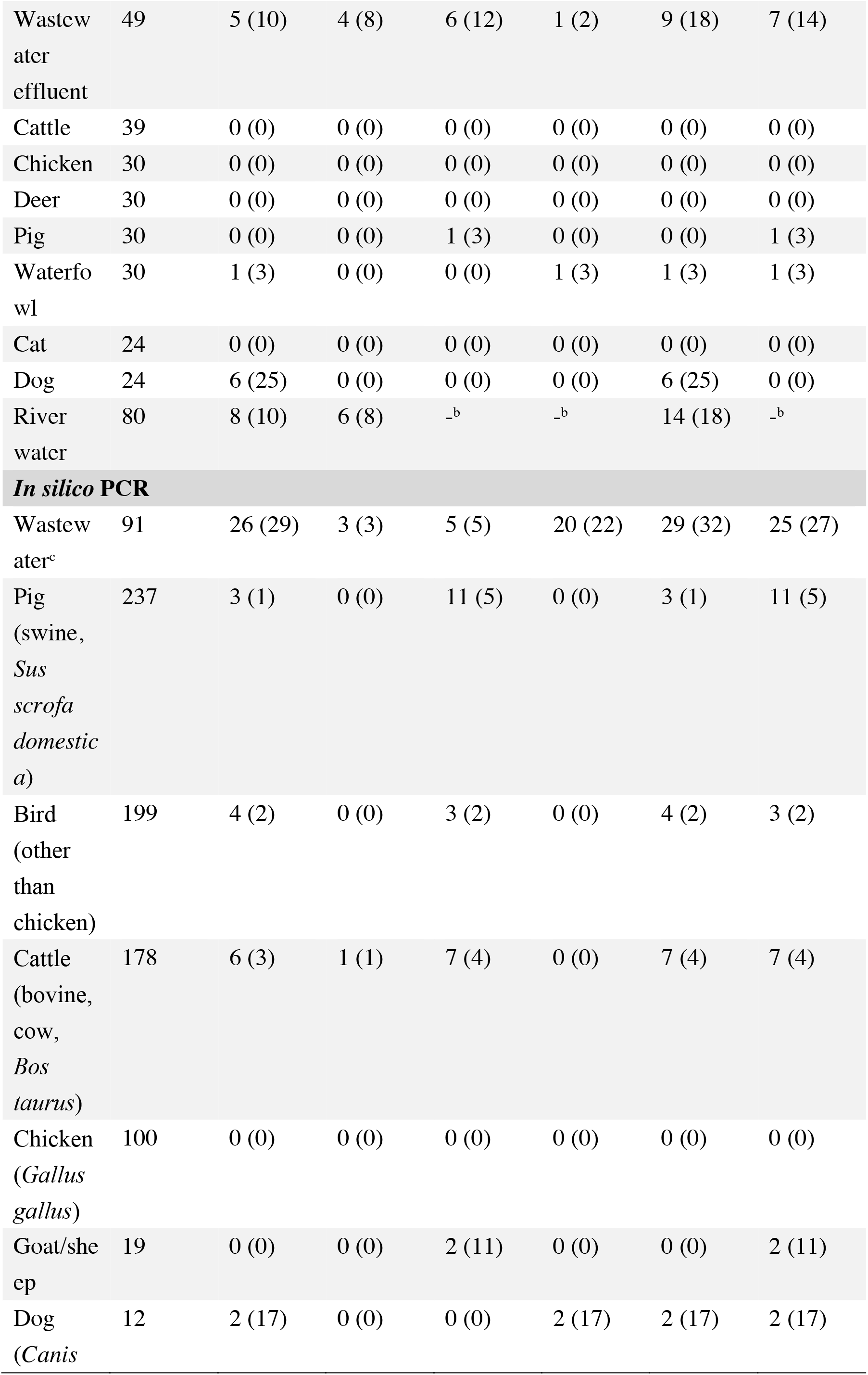
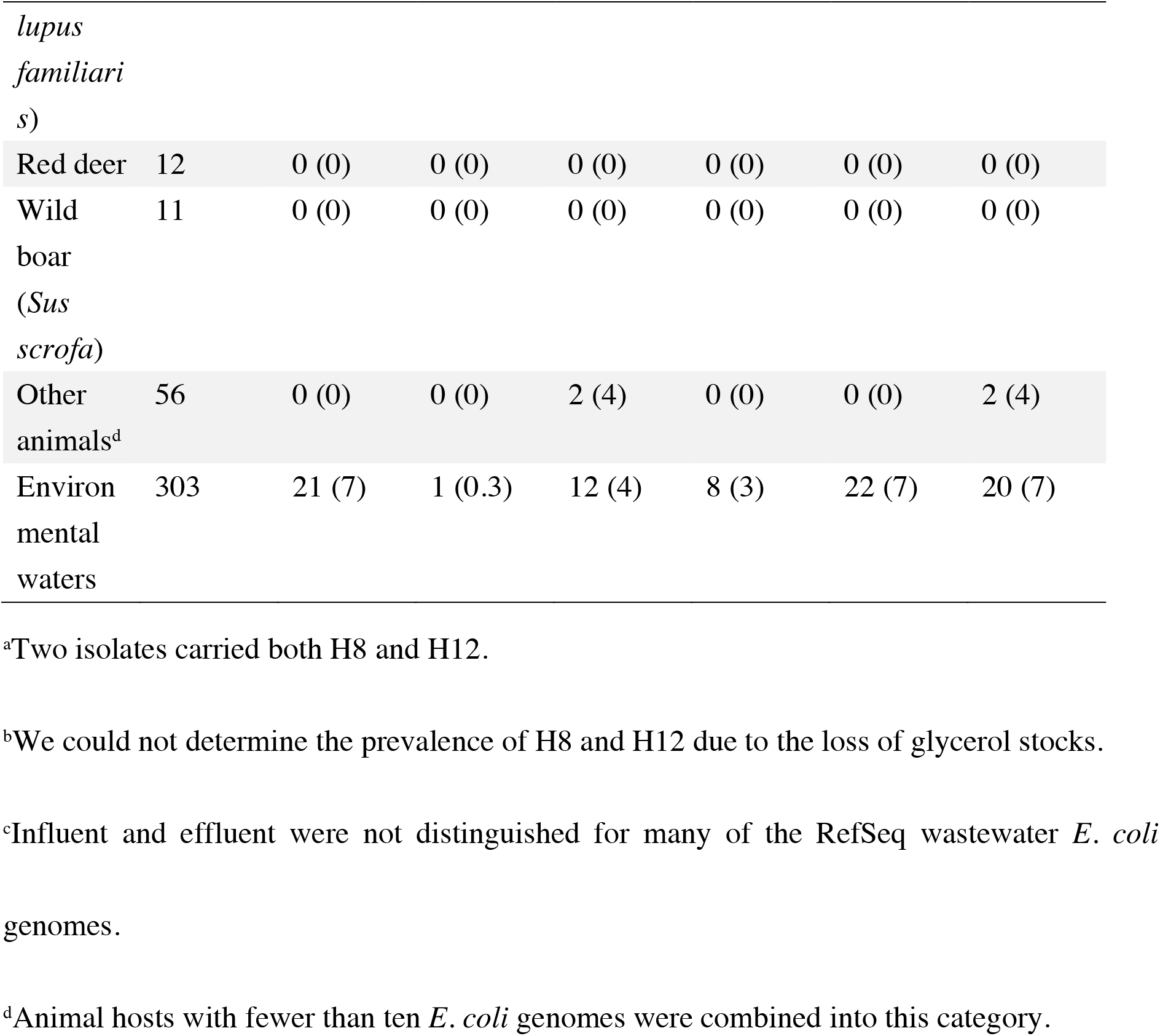
Prevalence of W_nqrC, W_clsA_2, H8, and H12 in *E. coli* determined by (*in silico*) PCR.

We also performed *in silico* PCR analysis using RefSeq *E. coli* genomes that were not used for blastn analysis. Among the 91 wastewater *E. coli* genomes, 29 genomes (31.9%) were positive for W_nqrC or W_clsA_2 (**Table 2, Table S9**). On the other hand, only 16 genomes (1.9%, seven from cattle, four from birds, three from pigs, and two from dogs) among the 824 animal *E. coli* genomes were positive for W_nqrC or W_clsA_2 (**Table 2, Table S10**). *In silico* PCR analysis of H8 and H12 revealed 25 wastewater *E. coli* genomes (27.5%) to be positive for H8 or H12, while 27 animal *E. coli* genomes (3.3%, 11 from pigs, seven from cattle, three from birds, two from goats, two from dogs, one from capybaras, and one from a South American tapir) were also positive for H8 or H12 (**Table 2, Table S9, Table S10**). In the *in silico* PCR analysis, the sensitivity and specificity of the W_nqrC and W_clsA_2 combination were higher than those of the H8 and H12 combination (31.9% vs 27.5% and 98.1% vs 96.7%).

### 3.7. Prevalence of MST markers in *E. coli* isolates from environmental waters

We performed PCR to determine the prevalence of W_nqrC and W_clsA_2 in *E. coli* isolates in river water. Of the 80 isolates tested, eight (10.0%) and six (7.5%) were positive for W_nqrC and W_clsA_2, respectively (**Table 2**). The prevalence of W_nqrC and W_clsA_2 was lower than but almost the same as that in *E. coli* isolates obtained from wastewater effluents (10.2% and 8.2%), which is congruent with the fact that the sampling points were strongly affected by wastewater.

We also performed *in silico* PCR on RefSeq environmental *E. coli* genomes. Of the 303 genomes, 21 genomes (6.9%), one genome (0.3%), 12 genomes (4.0%), and eight genomes (2.6%) were positive for W_nqrC, W_clsA_2, H8, and H12, respectively (**Table 2, Table S11**). The prevalence of MST markers was lower than in the RefSeq wastewater *E. coli* genomes: this is consistent with the fact that not all environmental *E. coli* originate from wastewater.

We note that the two markers developed in the present study, especially W_nqrC, are commonly found on plasmids and could be subjected to horizontal gene transfer (HGT) in the environment. Previous studies suggested that HGT could occur, for example, in biofilms, although HGT by conjugation rarely occurs between motile planktonic cells (Abe et al., 2020). Though we assume the contribution of HGT to the spread of the developed markers to be low in environmental waters, where water is constantly flowing, this should be kept in mind.

### 3.8. Concentrations of MST markers in wastewater samples

Concentrations of W_nqrC, W_clsA_2, H8, H12, and *uidA* in wastewater samples were determined by qPCR experiments (**Table S12**). The amplification efficiencies of qPCR assays were between 80%–100%, and the *r*^2^ values ranged from 0.988–0.999. The concentrations of markers and *uidA* in three primary effluent samples were: 7.13–7.24 log_10_ gene copies (GC)/L for W_nqrC; 8.39–8.54 log_10_ GC/L for W_clsA_2; 6.26–6.47 log_10_ GC/L for H8; 6.03–6.41 log_10_ GC/L for H12; and 7.21–7.32 log_10_ GC/L for *uidA*. The concentrations in the three biologically treated wastewater samples were: 4.43–4.76 log_10_ GC/L for W_nqrC; 6.12–6.18 log_10_ GC/L for W_clsA_2; 4.01–4.17 log_10_ GC/L for H8; 3.93–4.16 log_10_ GC/L for H12; and 4.50–4.80 log_10_ GC/L for *uidA*. Previously, *Bacteroides* HF183 and crAssphage markers were suggested to be two of the most promising sewage-associated MST markers (Ahmed et al., 2019a; Ahmed et al., 2019c). The mean concentrations of HF183 were previously reported to be 9.23 log_10_ GC/L for raw sewage and 7.32 log_10_ GC/L for treated effluent (Ahmed et al., 2016). The concentrations of crAssphage were previously reported to be 7.4–10.0 log_10_ GC/L for untreated wastewater, and the mean concentration was reported to be 6.5 log_10_ GC/L for treated effluent in high-income countries (Sabar et al., 2022). Although marker concentrations in wastewater vary by region and treatment system, *E. coli* MST markers appear to be less abundant in wastewater samples than markers such as HF183 and crAssphage.

The concentrations of W_nqrC were almost the same as *uidA* in all six samples, and the concentrations of W_clsA_2 were more than ten times higher than *uidA* in all the samples. This indicates that qPCR detected these marker genes not only from *E. coli* but also from non-*E. coli* species. *In silico* PCR analysis using the nr database on the Primer-BLAST website, which also includes the genomes of non-*E. coli* species, returned PCR products from non-*E. coli* species for both W_nqrC (e.g., *Klebsiella pneumoniae* and *Salmonella enterica*) and W_clsA_2 (e.g., *Pseudomonas aeruginosa* and *K. pneumoniae*). In the case of W_nqrC, 517 (93%) of 556 products were from *E. coli*, and 39 (7%) were from other species. In the case of W_clsA_2, only 29 (18%) of 157 products were from *E. coli*. Although detection of these marker genes in non-*E. coli* species does not preclude the applicability of W_nqrC and W_clsA_2 to a culture-independent method, further studies are needed to assess the applicability of these markers to culture-independent testing. We note that we also observed H8 to be prevalent among non-*E. coli* bacteria (Gomi et al., 2014), but it was proven in a subsequent study to be applicable in a culture-independent manner (Senkbeil et al., 2019).

### 3.9. Study limitations

This study has some limitations. First, when used in combination, the developed markers showed quite high specificity but showed relatively low sensitivity, indicating that a relatively large number of *E. coli* isolates would be needed in MST for each environmental sample. This was due to the tradeoff between the sensitivity and specificity of genetic markers (**Figure 2**). We assume this limitation to stem from using *E. coli* as an MST target organism rather than the methodologies employed in our study. A previous study attempted to classify *E. coli* genomes into different sources based on pan-genome information (Lupolova et al., 2017). The study analyzed a pan-genome of *E. coli* from different sources (avian, bovine, canine, environmental, human, and swine) and identified > 90,000 genes, three times more than the total genes identified in the present study. Although *E. coli* genomes could in many cases be classified into the correct sources, the classification was based on whole-genome sequence information, making it not ideal for MST applications. They also found a less clear association by host for *E. coli* when they compared the accessory genome trees of *E. coli* and another bacterial species, *S. enterica*. Moreover, there were subsets of animal isolates sharing significant genetic content with human isolates. The presence of the *E. coli* isolates that can colonize multiple host species is also reported in other studies (Johnson and Clabots, 2006; Li et al., 2019; Stenske et al., 2009). These factors indicate that *E. coli* is not the most suitable organism for MST studies. This is reflected in the fact that the present study found no MST markers with both high sensitivity and high specificity. However, we note that the sensitivity calculated in this study is at the isolate level, and sample-level sensitivity (i.e., sensitivity calculated for a wastewater sample, which can contain multiple *E. coli* strains) should be higher (Senkbeil et al., 2019). Moreover, the development of a multiplex PCR assay for detecting W_nqrC and W_clsA_2 in the same reaction will halve the time and labor needed to screen a large number of isolates.

Second, an in-depth analysis of the sensitivity and specificity of the developed markers was performed at the isolate/genome level but not in a culture-independent manner. Although we determined the concentrations of the markers in wastewater samples by qPCR, further studies are needed to assess the applicability (especially the specificity) of these markers in a culture-independent manner. This can be done by performing qPCR on DNA directly extracted from feces from different sources or by analyzing fecal metagenomes. In the latter case, one can estimate the abundance of the markers in each metagenome, for example, by read mapping. Analysis of metagenomes will remove the regional bias if metagenomes from geographically diverse hosts are used. Although further studies are needed to be able to use these markers in a culture-independent manner, we believe the two markers to be useful, since culture-dependent methods are used to detect *E. coli* in routine water quality assessments, and there is a need to identify the sources of the detected *E. coli* colonies. Moreover, cell suspensions prepared from

*E. coli* collected in routine water quality assessments can be used directly for PCR reactions without DNA extraction. We also found it possible to use colored *E. coli* colonies on selective media, in our case XM-G agar, for PCR reactions.

## 4. Conclusions

In the present study we identified two *E. coli* MST markers, namely W_nqrC and W_clsA_2, which can be used for tracking wastewater contamination in environmental waters. These two markers showed better performance than the previously developed H8 and H12 markers at the isolate level, but users should keep in mind that the sensitivity remained low, at around 20%– 30%, when used in combination. Although further studies are needed to test the applicability of the developed markers in a culture-independent manner, this study showed that the developed MST markers could be useful for identifying the sources of *E. coli* (whether they originated from domestic wastewater or not) at least at the isolate level. We believe that the PCR assays developed in this study can be readily applied to identify the sources of *E. coli* isolates collected during routine water quality assessments.

## Acknowledgements

This work was supported by the Environment Research and Technology Development Fund (JPMEERF20205006) of the Ministry of the Environment Japan, and Keihanshin Consortium for Fostering the Next Generation of Global Leaders in Research (K-CONNEX), established by Human Resource Development Program for Science and Technology, MEXT. Computations were partially performed on the NIG supercomputer at ROIS National Institute of Genetics. We thank those who provided animal fecal samples for this study.

## Supporting information

Supplementary information associated with this article can be found, in the online version, at

