## Supplementary Figures (S1-S5) for "Development of two microbial source tracking markers for detection of wastewater-associated *Escherichia coli* isolates"

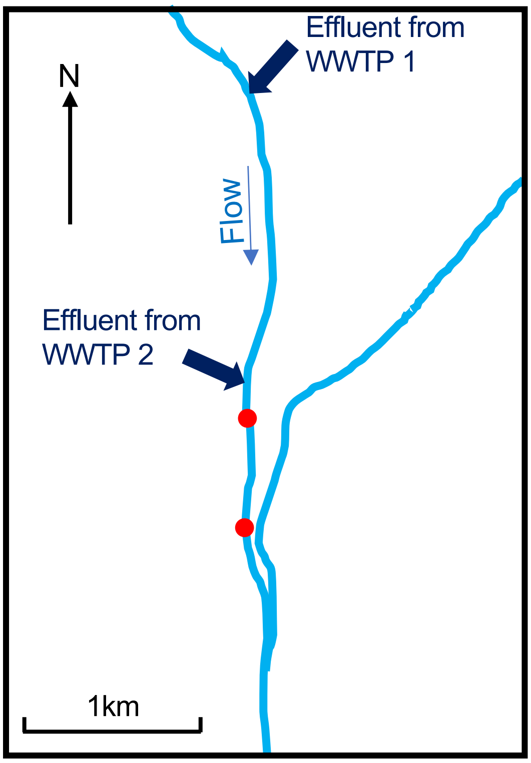


**Figure S1**. Map of sampling sites. Two red filled circles in the river indicate sampling sites. Four samples were collected from each sampling site.

(A) Cattle (n = 1446, including *Bos taurus*, bovine, beef cattle, cattle, calf, cow)

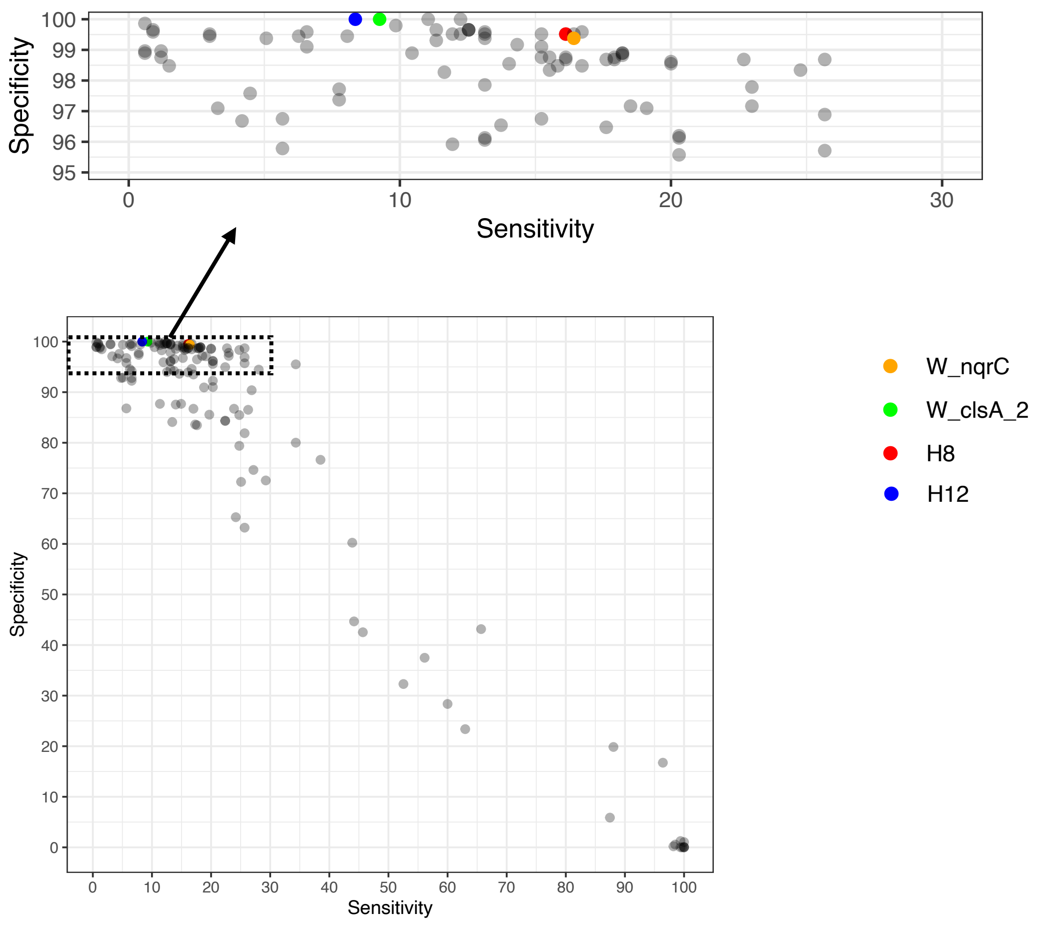

(B) Swine (n = 444, including pig, *Sus scrofa domesticus*, *Sus scrofa* (pig), swine, porcine)

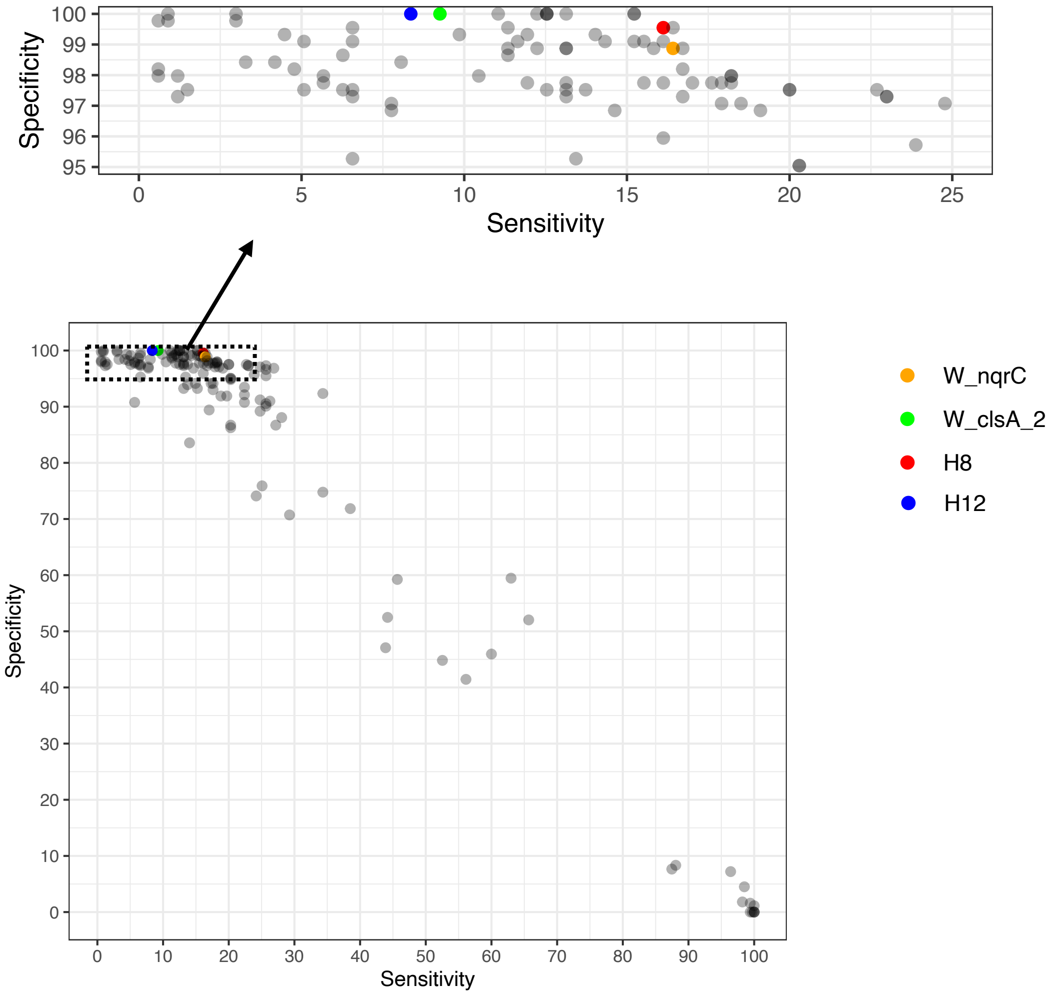

(C) Chicken (n = 302, including chicken, *Gallus gallus domesticus*, *Gallus gallus* (chicken))

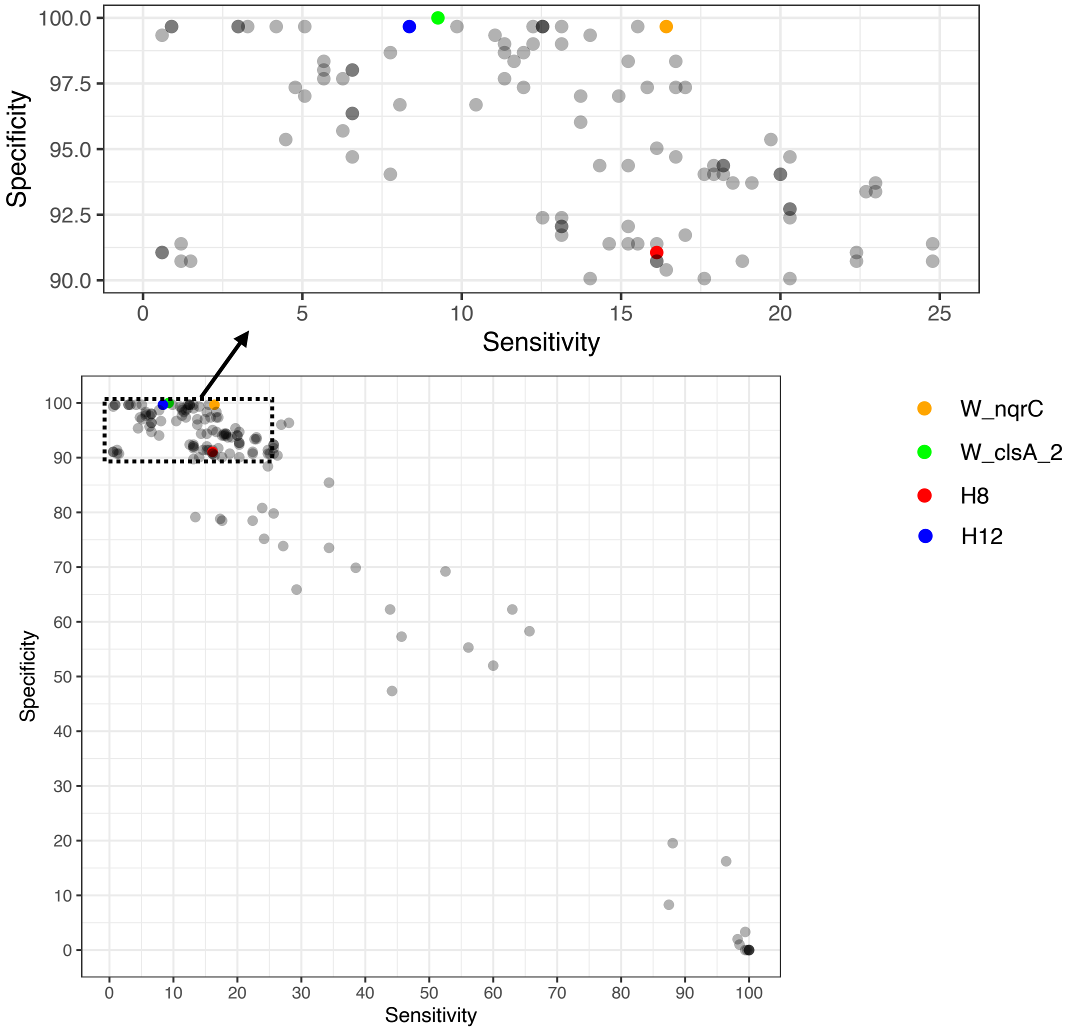

(D) Wild boar (*Sus scrofa*) (n = 186, including *Sus scrofa*, *Sus scrofa scrofa*)

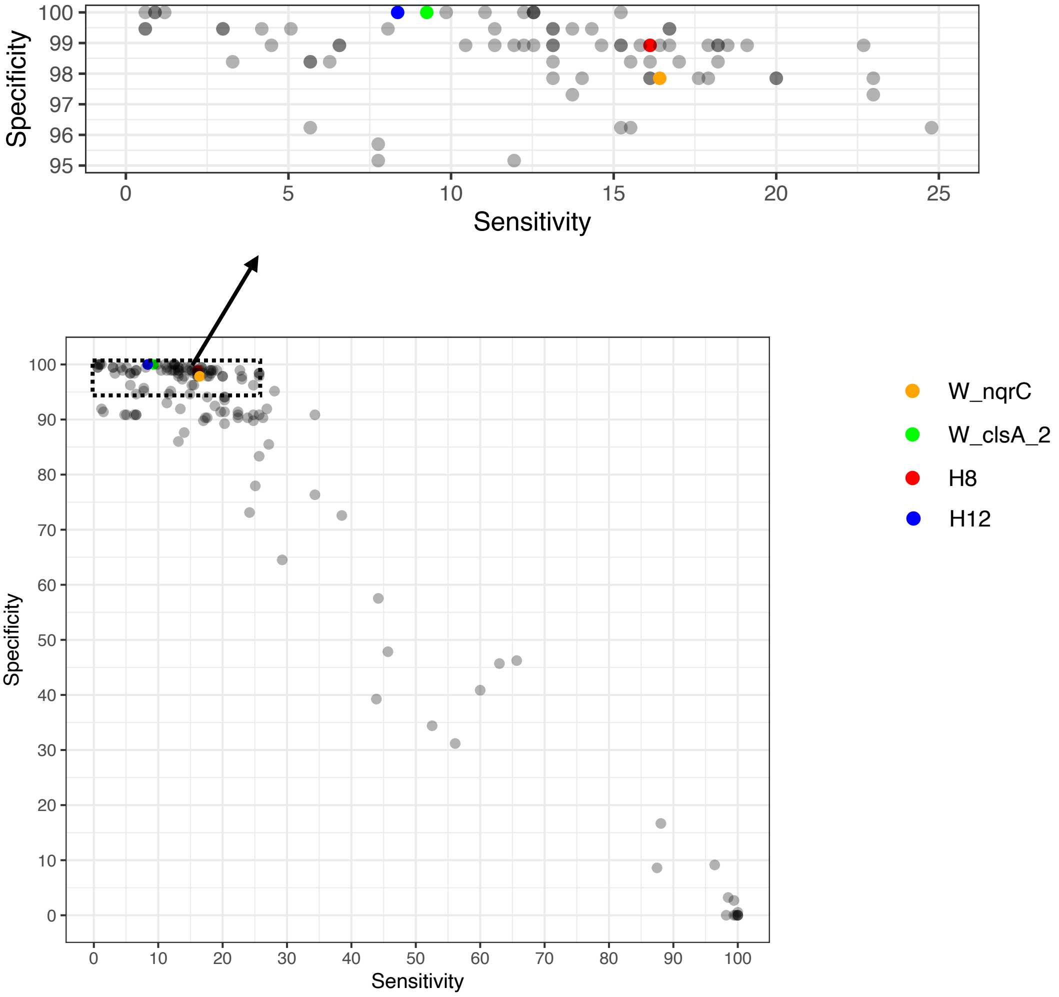

(E) Canine (n = 166, including *Canis lupus familiaris*, dog, canine)

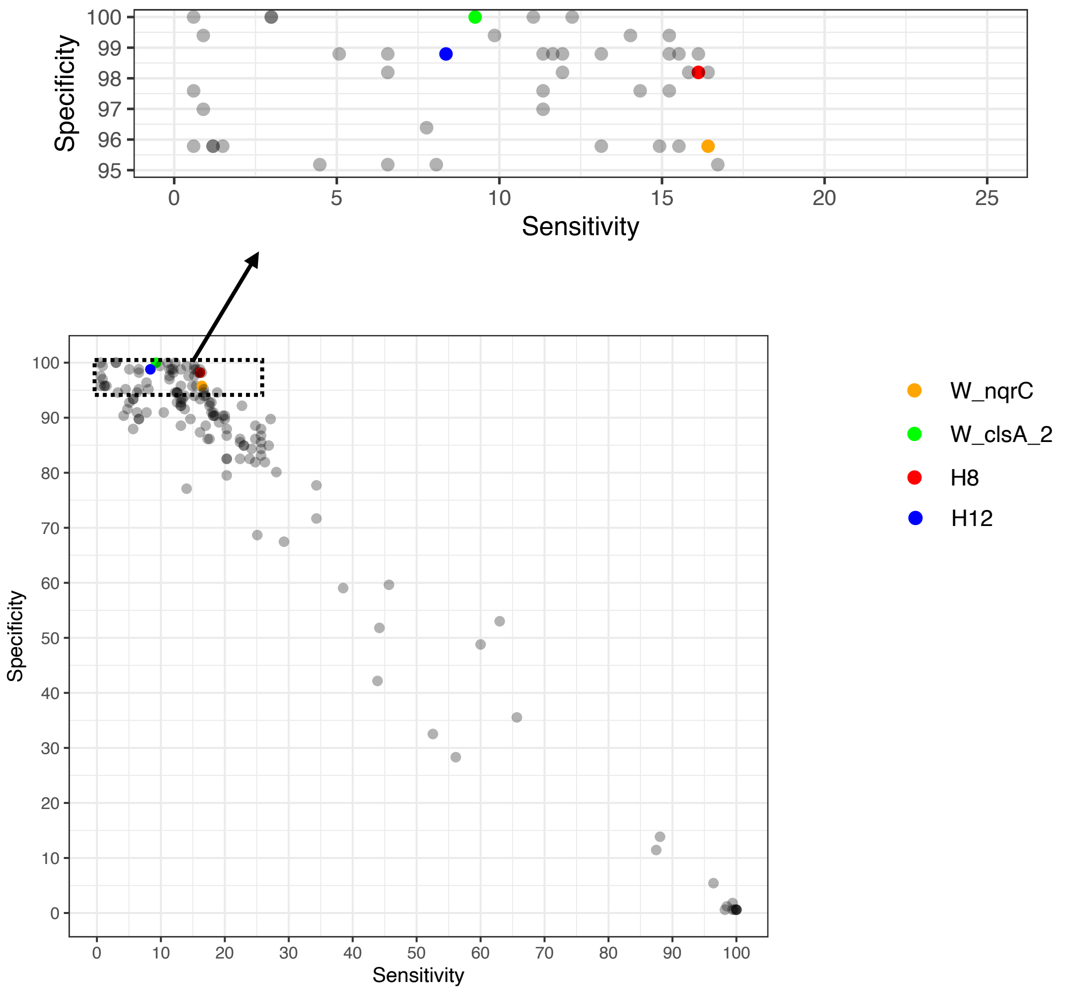

(F) Sheep and goat (n = 148, including *Ovis aries*, sheep, goat, ovine, capra)

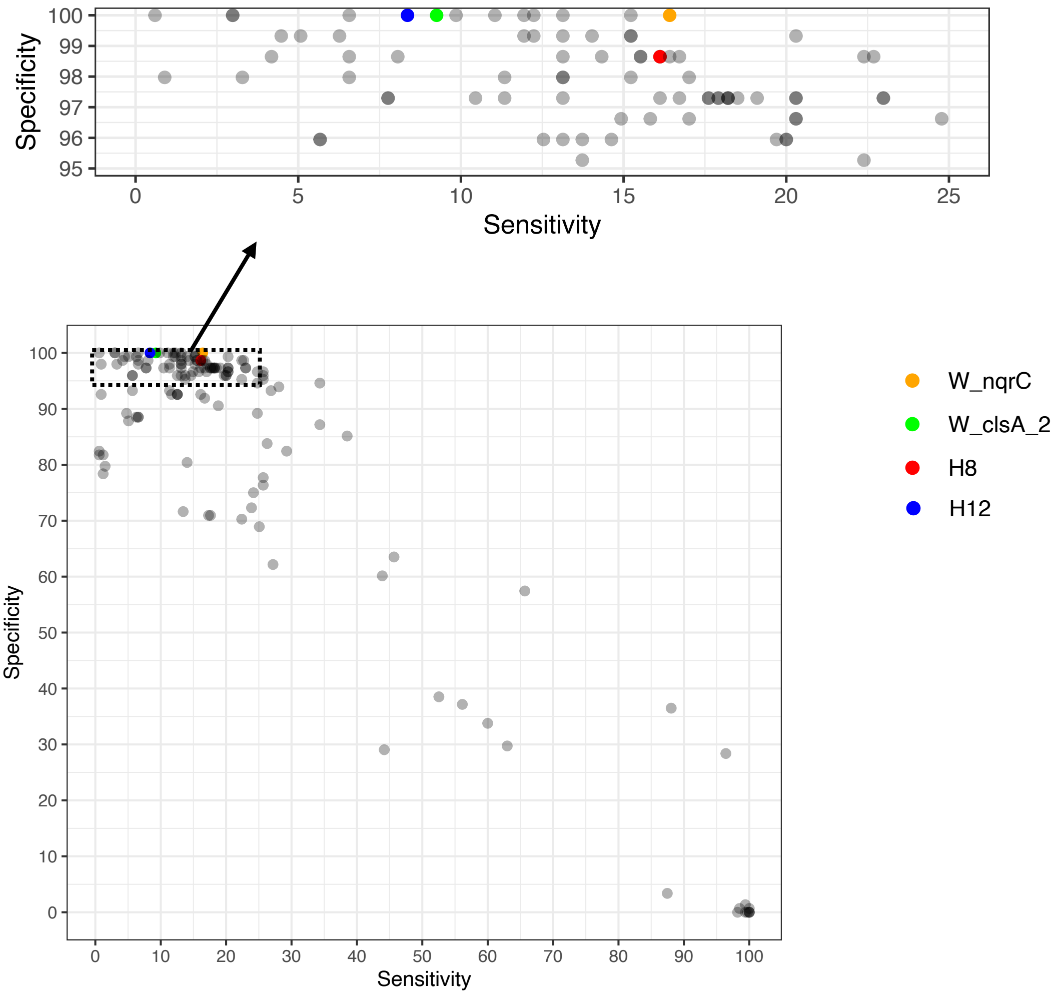

(G) Others (n = 626)

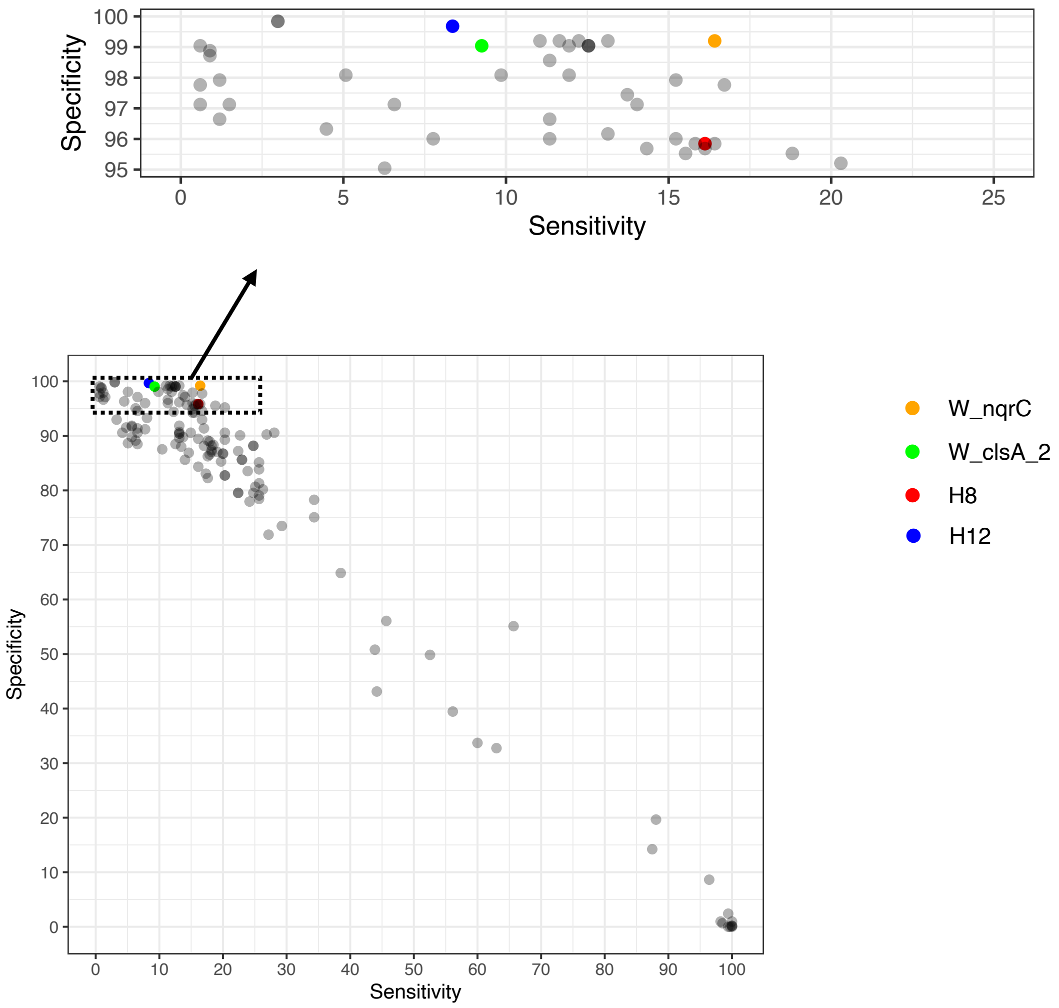


**Figure S2**. Sensitivity and specificity of candidate wastewater-associated marker genes shown separately for different host groups. Specificity was calculated using *E. coli* genomes from (A) "Cattle" (n = 1446, including *Bos taurus*, bovine, beef cattle, cattle, calf, cow), (B) "Swine" (n = 444, including pig, *Sus scrofa domesticus*, *Sus scrofa* (pig), swine, porcine), (C) "Chicken" (n = 302, including chicken, *Gallus gallus domesticus*, *Gallus gallus* (chicken)), (D) "Wild boar (*Sus scrofa*)" (n = 186, including *Sus scrofa*, *Sus scrofa scrofa*), (E) "Canine" (n = 166, including *Canis lupus familiaris*, dog, canine), (F) "Sheep and goat" (n = 148, including *Ovis aries*, sheep, goat, ovine, capra), (G) "Others" (n = 626). Host species with fewer than 100 *E. coli* genomes were combined into "Others".

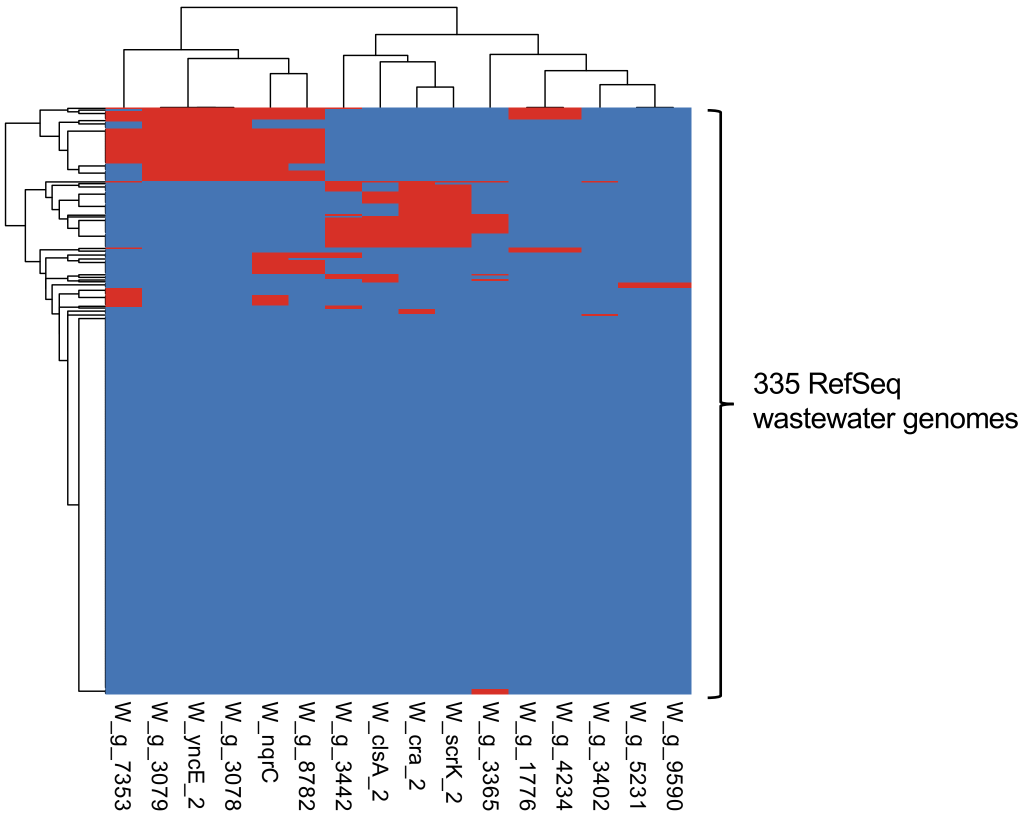


**Figure S3**. Heatmap of gene presence (red) and absence (blue) in 335 RefSeq wastewater genomes. Only candidate markers with >99% specificity are shown. Hierarchical clustering analysis was performed using the R package "pheatmap" (v1.0.12) with the "ward.D2" method.

(a)

>W_nqrC

ATGCGCAAGGGTAAAATTATTGTTGCATCAATGATGGTGCTTTGTGTCCTTATTTTT
GTTGCGGCGGCGGCATGGTTTATGCTGTTCCAGGGAGAAATGACGGAGCCGACAG
AGGAGGAAAAACAGGCGGCAATTTTACACGCTGCGGGCCTGATGAAATCAGAGA
CCCAGGATAAAAAATCAGTCGAAACGCTCTATCATCGTTATATTATTCAACGCCATG
TTAATTTAGATTCCGGCGAACTGGTGGCTGGCAGCAGCGCGGACACCGCGCGGCA
GAAGTGTGAAAAACTTGCCCCTGAACGCGACCCAGCGCAGGTTCGTCAGCGCTG
TACGGTGGCGGATGTGTTCTTTGTGAAAGATAAAAATAATGAAATTCAGCAGGTTA
TCATTCCAGTTACCGGTAAGGGCGCGAAATCCATGATG**CATGCGTTTCTTGCTCT**
**GGG**GCTGGACGGTCGTACGGTGAGAAATCTGTATTATTATCAGCAACGGGAAACG
CCGTTTCTGGGGG**CTAGGGTGGAAGACGCCAAC**TGGCGCAAACAGTGGCCAGG
GAAACGATTATTAGATAACAGCGGACATCCTGCATTGAAAATAGTACAGGATAAAC
CTGAACATGCCGATGAATATACCGTTGACGGTATTTCCGGTGCGACGTTGACATCA
ACCGGCGTTGAGAAAAGTATTAACTACTGGATGGGGCCGCAGGGATATGGTCAAT
TCTTACAGCGTCTTGCCAGCGATCGAAATAACCTTAATCTTTGA

(b)

>W_clsA_2

GTGTCCGACCCCAATGGGCTGACATGGGCAGCCACCCTCGTGTCGCTGGCGGTCG
CTATCCCCACAGCGGGGCATGCGGTCATCTACAAGCGTGACCCTCGATCGGCCAC
GCTGTGGGTACTGCTGATCGCCTTGCTGCCGCTGGGCGGTTCGCTGCTGTATGGGC
TGTTCGGCATCAACCGTTACCAGCGGCGGGCGCGGCGGCTCTTCCCGGGGGCAGA
TCCTGCTGTTCGGCAGGATTTCTCGCCCACGATACCCGAGGCTGTATCGCCGCCAC
TTGCCGGTCTGGCCCACCTGGTGGGACGCGCCACCGGCCAGTCGCTGACCGGCG
GCAACCGCATCGAGCCGCTTGTCGATGGCGAGCAGGCCTACCCGGCCATGCTCGC
CGCCATCGAATCGGCGCGGTACAGCGTCGCCTTGACCTCGTACATCTTCGACAGCC
AAGGCATCGGGGCGCAGTTCGTCGATGCGCTGCGCCGGGCTCATGAGCGCGGCGT
GCAAGTGCGGGTGCTGATCGACGACGTGTATGCCCGCTGGACGCCCCGCAGCGCC
TACCGCGCCTTGCAGCGCGCCGGCGTTCCGGCGGCGACGTTCAACCCGACATTGA
TTCCCGCGCGCCTGCATGCCGCGCATCTGCGGAATCACCGCAAGCTGCTGGTGAT
CGATGGCGAGACGGGTTTCACCGGCGGATTGAACATCTTCAGCCCGTACTGGCGG
CCCGATGCGCCGGACCAGGCCTGTCACGATTTGCACTTTCGTCTGCGCGGGCCGG
TGGTGGAGCATCTGATGCGGAGCTTC**ACCGATGATTGGTGCGATAC**CACGGGCG
AGCGGCTCAACGAGGGTTTCTGGGGCGAACCGCCCGTAACGGCTGATGAGCCGG
GAACCAGTTGGGCGCGCGGCATCGAGGCCGGTCCCGACGAGGCCCTGGACCGGA
TGCGCTGGAC**CTTCATGGGCGCTTTGAGC**GCGGCGAAGCATTCGGTGCGCATCT
GGACACCTTACTTCGTCCCCGATCAGCCGATGATCGCGGCGCTGAGCACGGCCGC

GTTGCGCGGCGTGCGTATCGACGTGCTGACGCCCGCAAACGGCGACCATCCCACG
GTGCAATGGGCCGCGCGCGCCCACTACTGGCAGGTGCTGGAGCACGGTGTGCGC
ATCTTCGAACGGCCTGGCCCGTTCGACCACAGCAAGCTGATGCTGGTAGACGGAG
CGTGGTGCTGCCTGGGTTCGGCCAATTGGGATGCGCGCAGTTTGCGCCTCAACTT
CGAGTTCAATGTGGAGGTGTACGACACCGCGTTGTCTACGCGGCTGTCATCCCTTT
TTGATGCCGCTCGTGATGCGTCGGGCGAGATATCGGCGAAGGCGTTGCGCGCCCA
CCCGCTGGCCATGCGCTTGCGCGACGGGGTGGCCCGTCTGTTCACCCCCATTCTCT
AG

**Figure S4**. The sequences of (a) W_nqrC and (b) W_clsA_2. Primer binding sites are underlined.

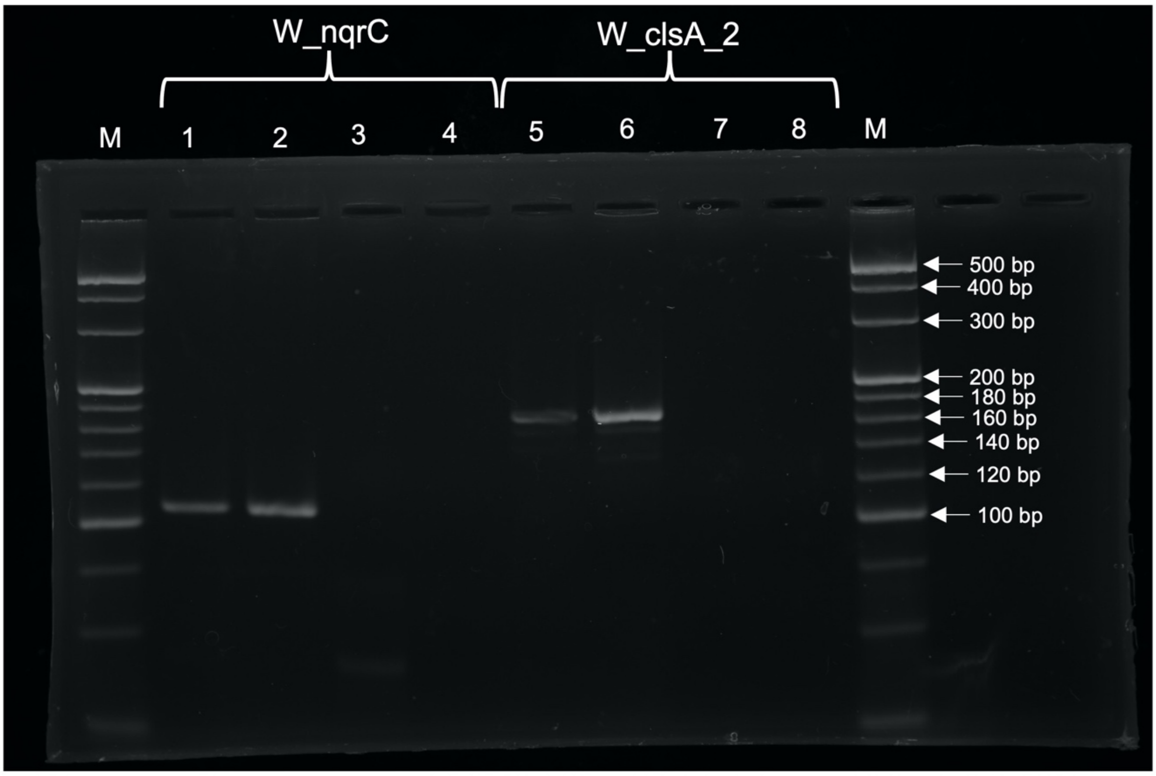


**Figure S5**. Agarose gel electrophoresis of the *E. coli* MST PCR amplicons. Lanes: M, DNA ladder; 1, KMi027 (positive control); 2, positive isolate obtained from wastewater; 3, KOr026 (negative control); 4, water (negative control); 5, KOr026 (positive control); 6, positive isolate obtained from wastewater; 7, KMi027 (negative control); 8, water (negative control).

**Supplementary Tables**

See the excel file for the following tables:

**Table S1**. Basic information on 50 wastewater *E. coli* genomes sequenced by us.

**Table S2**. Basic information on 82 complete RefSeq animal *E. coli* genomes.

**Table S3**. Basic information on 335 RefSeq wastewater *E. coli* genomes used for blastn analysis.

**Table S4**. Basic information on 3318 RefSeq animal *E. coli* genomes used for blastn analysis.

**Table S5**. Basic information on 414 *E. coli* isolates used for PCR.

**Table S6**. Candidate wastewater-associated marker genes identified by Scoary.

**Table S7**. Characteristics of *E. coli* genomes positive for W_nqrC.

**Table S8**. Characteristics of *E. coli* genomes positive for W_clsA_2.

**Table S9**. Basic information on 91 RefSeq wastewater *E. coli* genomes used for *in silico* PCR.

**Table S10**. Basic information on 824 RefSeq animal *E. coli* genomes used for *in silico* PCR.

**Table S11**. Basic information on 303 RefSeq environmental *E. coli* genomes used for *in silico* PCR.

**Table S12**. Concentrations of genetic markers in wastewater samples determined by qPCR.
